# Learning the human chromatin network from all ENCODE ChIP-seq data

**DOI:** 10.1101/023911

**Authors:** Scott M. Lundberg, William B. Tu, Brian Raught, Linda Z. Penn, Michael M. Hoffman, Su-In Lee

## Abstract

**Introduction**: A cell’s epigenome arises from interactions among regulatory factors — transcription factors, histone modifications, and other DNA-associated proteins — co-localized at particular genomic regions. Identifying the network of interactions among regulatory factors, the *chromatin network*, is of paramount importance in understanding epigenome regulation.

**Methods**: We developed a novel computational approach, ChromNet, to infer the chromatin network from a set of ChIP-seq datasets. ChromNet has four key features that enable its use on large collections of ChIP-seq data. First, rather than using pairwise co-localization of factors along the genome, ChromNet identifies *conditional dependence* relationships that better discriminate direct and indirect interactions. Second, our novel statistical technique, the *group graphical model*, improves inference of conditional dependence on highly correlated datasets. Such datasets are common because some transcription factors form a complex and the same transcription factor is often assayed in different laboratories or cell types. Third, ChromNet’s computationally efficient method and the group graphical model enable the learning of a joint network across all cell types, which greatly increases the scope of possible interactions. We have shown that this results in a significantly higher fold enrichment for validated protein interactions. Fourth, ChromNet provides an efficient way to identify the genomic context that drives a particular network edge, which provides a more comprehensive understanding of regulatory factor interactions.

**Results**: We applied ChromNet to all available ChIP-seq data from the ENCODE Project, consisting of 1451 ChIP-seq datasets, which revealed previously known physical interactions better than alternative approaches. ChromNet also identified previously unreported regulatory factor interactions. We experimentally validated one of these interactions, between the MYC and HCFC1 transcription factors.

**Discussion**: ChromNet provides a useful tool for understanding the interactions among regulatory factors and identifying novel interactions. We have provided an interactive web-based visualization of the full ENCODE chromatin network and the ability to incorporate custom datasets at http://chromnet.cs.washington.edu.

## Introduction

Regulatory factors — such as transcription factors, histone modifications, and other DNA-associated proteins — co-localize in the genome and interact with each other to regulate gene expression [15], the physical structure of the genome [10], cell differentiation [5], and other cellular processes. Identifying the genomic co-localization in this network among regulatory factors, the *chromatin network*, is important for understanding genome regulation and the function of each regulatory factor [60, 4]. To identify the chromatin network, we can use chromatin immunoprecipitation-sequencing (ChIP-seq) to measure genome-wide localization of regulatory factors, and then compare ChIP-seq datasets to find regulatory factors that co-localize [46, 11]. Co-localization may indicate that two factors interact physically, by forming a complex, or functionally, such as by regulating similar DNA targets.

However, identifying pairwise co-localization alone fails to distinguish direct interactions from indirect interactions. A direct interaction represents physical contact or close functional coupling that requires spatial proximity. An indirect interaction is not from physical contact or direct functional coupling, but instead reflects the transitive effect of other direct interactions. Consider a simulated chromatin network among four factors, where factor C recruits A and B, and A in turn recruits D (Figure 1A, top). Because all pairs of ChIP-seq datasets are correlated to each other (Figure 1A, middle), a simple co-localization method would incorrectly infer interactions among all the factors (Figure 1A, bottom left). In a *conditional dependence network* (Figure 1A, bottom right), if two variables (here, factors) are *conditionally dependent*, then there is an edge between them. The *conditional dependence* between two factors measures their co-localization after accounting for information provided by other factors. If we infer a conditional dependence network, we eliminate indirect edges from the network, such as between factors A and B, because their co-localization at peaks 3 and 5 can be *explained away* by another factor C (C recruits A and B). Hence, incorporating more ChIP-seq datasets allows more indirect edges to be removed, resulting in a higher quality inferred network.

**Figure 1:**
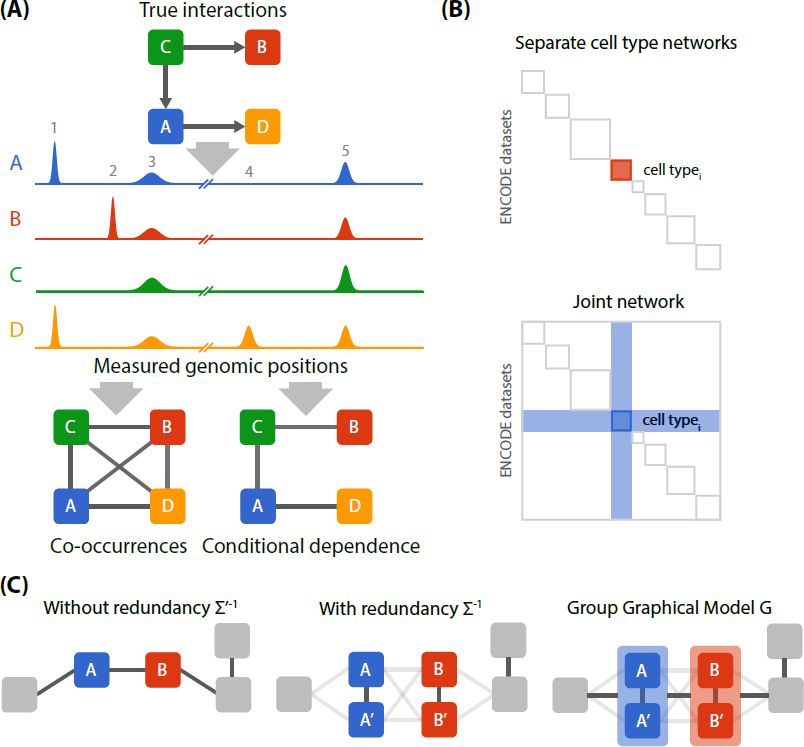
**(A)** *Top*: Interaction network among four simulated regulatory factors. *Middle*: Binding activity from simulated ChIP-seq datasets, where each peak represents a putative binding position of a protein. *Bottom*: Networks inferred from ChIP-seq datasets based on co-occurrence (left) or conditional dependence (right). **(B)** Comparison of separate cell type networks (top) with a single joint network (bottom). In a joint model, factors in each cell type have opportunities to be connected with new regulatory factors in other cell types, as highlighted by the blue shaded region (bottom). **(C)** Redundant information obscures conditional dependence connections. *Right*: Without redundancy, standard methods robustly infer a conditional dependence network. *Middle*: Highly correlated variables (such as *A* and *A’*) are strongly connected with each other and lose their connections with other variables. *Left*: A Group Graphical Model (GroupGM) represents the conditional dependence between groups of correlated variables, which restores the connection between *A* and *B*.

Here we present ChromNet, an approach that estimates the human chromatin network using a conditional dependence network among regulatory factors from 1,451 human ENCODE ChIP-seq datasets (Supplementary Table 1). Integrating all ENCODE datasets from many cell types into a single network provides several advantages. First, it enables the extraction of global patterns in the conditional dependence relationships among regulatory factors in all cell types. Second, it provides a flexible model that allows direct comparison of cell type specific sub-networks because factors are conditioned on the same global set of ChIP-seq datasets across all cell types. Finally, it greatly increases the number of edges to consider by allowing edges connected to factors outside a single cell type (Figure 1B). We show that this leads to a substantially increased enrichment fold for known protein interactions.

Learning this network involves three key challenges. First, learning a network among thousands of ChIP-seq datasets based on millions of genomic regions is highly computationally intensive. To solve this challenge, we utilized an efficient approach that involves the computation of an *inverse correlation matrix*, which does not require an expensive iterative learning procedure. This is in contrast to some other methods, such as Bayesian networks [62, 3] and Markov random fields [72], which face difficulties scaling up and make it infeasible to run on all 1,451 ChIP-seq datasets (Supplementary Note 1). Second, some regulatory factors are in the same complexes, and factors are often measured in different labs, conditions, or cell types, which creates significant correlations in the data. When some variables are highly correlated with each other, standard methods often learn edges only among these variables and disconnect them from the rest of the network (Figure 1B, middle) [2]. Incorporating more ChIP-seq datasets exacerbates this problem.

To solve this challenge, we present the *Group Graphical Model* (GroupGM) representation of a conditional dependence network that expresses conditional dependence relationships among groups of regulatory factors as well as individual factors (Figure 1B, bottom). We show that GroupGM improves the interpretation of a conditional dependence network by allowing edges to connect groups of variables, which makes the edges robust against data redundancy. Third, network edges can be driven by interactions in specific genomic contexts. To help understand these contexts we present an efficient method to label every genomic position with its impact on an inferred GroupGM edge.

Previous work on learning interactions among regulatory factors from ChIP-seq data used much smaller data collections. ENCODE identified conditional dependence relationships among groups of up to approximately 100 datasets in specific genomic contexts [21]. Other authors used partial correlation on 21 datasets [34], Bayesian networks for 38 datasets [36], or partial correlation and penalized regression on either 27 human datasets [54] or 139 mouse embryonic stem cell datasets [26]. Still other authors used a Markov random field with 73 datasets in *D. melanogaster* [72], a Boltzmann machine with 116 human transcription factors [44], or bootstrapped Bayesian networks in 112 regulatory factors in *D. melanogaster* [62, 3]. Only other approaches also based on linear dependence models, such as partial correlation used by Lasserre et al. [34], scale well to all ENCODE datasets (‘Partial correlation’ and ‘rank(Raw read pileup)’ in Supplementary Figure 1). The ChromNet approach extends these methods in four distinct ways: 1) We show that linear dependence models can directly be applied to the genome-wide untransformed read count data (Supplementary Figure 1); 2) ChromNet addresses a fundamental challenge in network estimation when some of the variables are highly correlated with each other (collinearity) through a novel statistical method, the group graphical model; 3) ChromNet uses a novel method to identify genomic positions and genomic contexts that drive specific network edges; and 4) Jointly modeling multiple cell types leads to a more informative network with a substantially higher enrichment for known protein interactions. Network inference has also been applied to gene expression data, but the number of available samples in expression data is much lower than for ChIP-seq datasets, which leads to different challenges than those encountered in regulatory factor networks [41].

ChromNet departs from previous approaches by enabling the inclusion of all 1,451 ENCODE ChIP-seq datasets into a single joint conditional dependence network. GroupGM and an efficient learning algorithm allow seamless integration of all datasets comprising 223 transcription factors and 14 histone marks from 105 cell types without requiring manual removal of potential redundancies (Supplementary Table 1). We show that this approach significantly increases the proportion of network relationships among ChIP-seq datasets supported by previously known protein-protein interactions as compared to other scalable methods (Results). We also demonstrate the potential of ChromNet to aid new discoveries by experimentally validating a novel interaction.

## Results

### Uniformly processed data reduces noise when learning conditional dependence

To ensure comparable signals across all ChIP-seq datasets, we re-processed raw ENCODE sequence data with a uniform pipeline (Figure 2A). We downloaded raw FASTQ files from the ENCODE Data Coordination Center [11, 56, 16] and mapped them using Bowtie2 [33] to the human genome reference assembly (build GRCh38/hg38) [20]. We binned mapped read start sites into 1,000–base-pair bins across the entire genome, which results in a 3,209,287 x 1,451 data matrix X where genomic positions are viewed as samples (Figure 2A). We compared several different data pre-processing methods and chose binned read counts for three reasons: 1) they allow easy integration of external ChIP-seq experiments, 2) they do not require the determination of various cut-offs in a peak calling algorithm, and 3) a network inferred from read counts performs well revealing previously known protein-protein interactions (Supplementary Figure 1).

**Figure 2:**
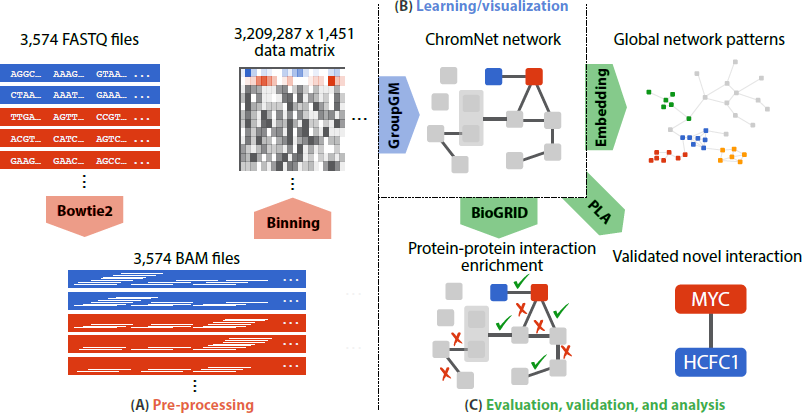
**(A)** Uniform processing pipeline includes aligning sequences using Bowtie2, and then binning them into 1,000-base-pair regions. **(B)** We inferred the GroupGM from all 1,451 ChIP-seq datasets and integrated the learned model into a web interface to facilitate broad use. **(C)** We evaluated the learned model against known physical protein interactions (BioGRID), mapped global patterns through network embedding, and validated a novel predicted MYC-HCFC1 interaction with a Proximity Ligation Assay (PLA).

### A conditional dependence network can be efficiently learned from binned read count data

Learning a conditional dependence network among thousands of ChIP-seq datasets each containing millions of samples (genomic positions) requires an efficient algorithm (Figure 2B). It is well known that the nonzero pattern of the *inverse covariance matrix* of Gaussian random variables represents the conditional dependence network [35, 40]. The inverse correlation matrix, 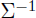, is just a normalized version of the inverse covariance matrix and also represents conditional dependence. A zero element 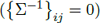 means that the *i*th and *j*th variables are conditionally independent of each other given all other variables—they are not connected by an edge.

While it is common practice to learn the conditional dependence network among continuous-valued variables based on the estimation of 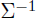 [24], count data requires more care. Distributions of counts in binned ChIP-seq reads are often clearly truncated at zero, and also increase in variance for high read counts. Multivariate distributions with count-valued marginal distributions are often very restrictive (for example only allowing positive correlations) or are infeasible to estimate for thousands of dimensions [68]. An often employed alternative is to use a multivariate Gaussian distribution after appropriately transforming the count data, such as with the *sqrt* or *asinh* function [9]. However, interestingly, our results show that applying a linear Gaussian model directly to the binned read counts of ENCODE ChIP-seq data better recovers known protein-protein interactions than when using standard normalizing data transforms (Methods; Supplementary Figure 1). This leads to an efficient and simple model formulation for ChromNet applied directly to the mapped read counts, which is relatively easier to obtain compared to other ChIP-seq data pre-processing methods and does not require any threshold.

ChromNet first computes the *inverse sample correlation matrix* 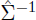 from the data matrix *X* of 1,451 variables and 3,209,287 samples, and then uses a GroupGM approach to interpret elements of 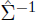 as weights of network edges (Figure 2B).

### Group modeling mitigates the effects of redundancy

Many ENCODE ChIP-seq datasets contain redundant positional information. Conventional conditional dependence methods have a key limitation in modeling redundant data. If datasets *A* and *A’* are highly correlated, a conventional method would connect *A* with *A’* but connect *A* to the rest of the network only weakly (Figure 1B). Arbitrarily removing or merging redundant datasets can hide or eliminate important information in the data.

GroupGM overcomes challenges with redundant data in conditional dependence models by allowing edges that connect groups of datasets (such as [*A,A*^’^] and [*B,B*^’^]). A group edge weight represents the total dependence between the variables in the two groups that the edge connects, and is computed from 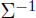 as (Figure 1C, Methods):

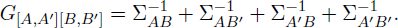

An edge in a GroupGM model implies conditional dependence between the linked groups, but does not specify the involvement of individual factors in each group. We prove that GroupGM correctly reveals conditional dependencies in the presence of redundancy (Supplementary Note 2).

A group is defined as a set of highly correlated variables whose individual conditional dependence relationships with other variables are not likely to be captured, as illustrated in Figure 1C. To obtain groups, we used complete linkage hierarchical clustering, and restricted groups to have minimum pairwise correlation of *ρ* (= 0.8) within each group. The choice of complete-linkage clustering allows us to obtain groups where all the factors are highly correlated. Because the complete-linkage distance metric merges two clusters based on the minimum correlation between any two variables in the groups, we can stop merging when the minimum correlation becomes less than or equal to *ρ* before creating all 2*p*–1 groups, where *p* = 1,451.

Each variable (a ChIP-seq dataset) can be in multiple groups as long as it is highly correlated with at least one dataset. This multi-scale nature of groups is a unique feature of the group graphical model. It allows us to capture multiple ways each factor can be connected with other factors. Say that a dataset for factor A forms a group with another dataset for factor B. In the group graphical model, A can have connections specific to itself and connections shared with B, and their edge weight values would indicate which connections are statistically robust. This allows us to reveal multiple kinds of interactions A can have – specifically to itself and to A and B as a complex. The later may not be captured by a conventional conditional dependence network, such as inverse correlation or partial correlation, if A and B are highly correlated with one another.

The purpose of having a threshold for minimum pairwise correlation ρ is to identify sets of variables whose high within-group correlation is likely to prevent them from being connected to other variables in the network. The threshold used in this paper *ρ* = 0.8 captures 53% of all the multi-factor groups formed by hierarchical clustering, and was chosen so as to include strong groups while still keeping the size of groups small enough to interpret (Supplementary Figure 2).

### Conditional dependence and joint group modeling improve the recovery of known protein-protein interactions

To evaluate how conditional dependence and group modeling both contribute to the performance of ChromNet we estimated three networks among ChIP-seq datasets using the following three methods, where each method produces a set of weighted edges:

1. **Correlation**: We learned a naive co-occurrence network, using pairwise Pearson’s correlation between all pairs of datasets.
2. **Inverse correlation**: We learned a conditional dependence network, by computing the inverse of the correlation matrix.
3. **GroupGM**: We learned a group conditional dependence network, which addresses tight correlation among datasets by allowing edges between groups of variables.

Partial correlation is similar to inverse correlation and performs nearly as well (Supplementary Figure 3). We did not include other previously described methods because they do not scale to the large data collection we used (Supplementary Note 1).

To assess the quality of the estimated networks, we identified the edges corresponding to published protein-protein interactions. As ground truth, we used the BioGRID database’s assessment of physical interactions between human proteins from experiments deemed low throughput [61]. For evaluation, we excluded edges connecting the same regulatory factor even when measured in different labs, cell types, or treatment conditions. These edges were excluded from evaluation to prevent them from artificially inflating the accuracy of the methods. We also excluded edges involving a histone mark because they do not exist in BioGRID. For these edges we ran a separate evaluation using the HIstome database [27] and showed that the group graphical model shows higher enrichment than the alternative methods (Supplementary Figure 4). When we measured the conditional dependence between a pair of ChIP-seq datasets in GroupGM, to avoid the inclusion of many redundant edges, for each pair of datasets, we picked the maximum edge weight out of all network edges connecting groups, each of which contains one of the corresponding datasets. This way, we consider exactly the same number of dataset pairs for evaluation across all methods. We only scored edges from groups containing a single type of factor (about half of the groups; see Supplementary Figure 2), because if a group contains more than one factor, there is no clear way to characterize such an edge as true or false from BioGRID, or match it with an edge from competing methods for comparison.

### Group modeling improves the recovery of interactions within and between cell types

We compared performance of the three methods described above across a range of prediction thresholds. For each network, we varied a number *N* of evaluated edges from 1 to the total number of edges. For each value of *N*, we identified the set of *N* edges with the largest weights. We also randomly picked *N* edges without regard to weight rank as a background set. We then calculated how many edges in each set matched known protein-protein interactions from BioGRID. We computed fold enrichment by dividing the number of matched edges in the prediction set by the expectation of the number in the background set. Since 8.4% of datasets pairs in the same cell type are supported by a BioGRID physical interaction, an enrichment fold of 1 corresponds to 8.4% of recovered edges matching prior knowledge. Enrichment fold captures the effect of both Type I and Type II error rates (see Methods).

We first measured performance within all cell types, excluding edges between datasets in different cell types (Figure 3A top). Since the limited number of annotations in BioGRID imperfectly represent the human chromatin network, one cannot draw strong conclusions about absolute performance from this benchmark. Relative performance of the methods, however, is clear. Inverse correlation performed better than correlation, and GroupGM outperformed inverse correlation. This supports the idea that better resolution of direct versus indirect interactions contributes to improved performance of inverse correlation over correlation, while greater robustness against relationship-hiding redundancy likely contributes to improved performance of GroupGM over inverse correlation. The value of conditional dependence and group modeling is also further supported by specific examples in the network (Figure 4; Supplementary Figure 5; Supplementary Figure 6), and by the fact that Group GM still outperforms inverse correlation even after attempting to remove the strongest redundancies by merging datasets from different labs targeting the same factor in the same cell type/condition (Supplementary Figure 7).

**Figure 3:**
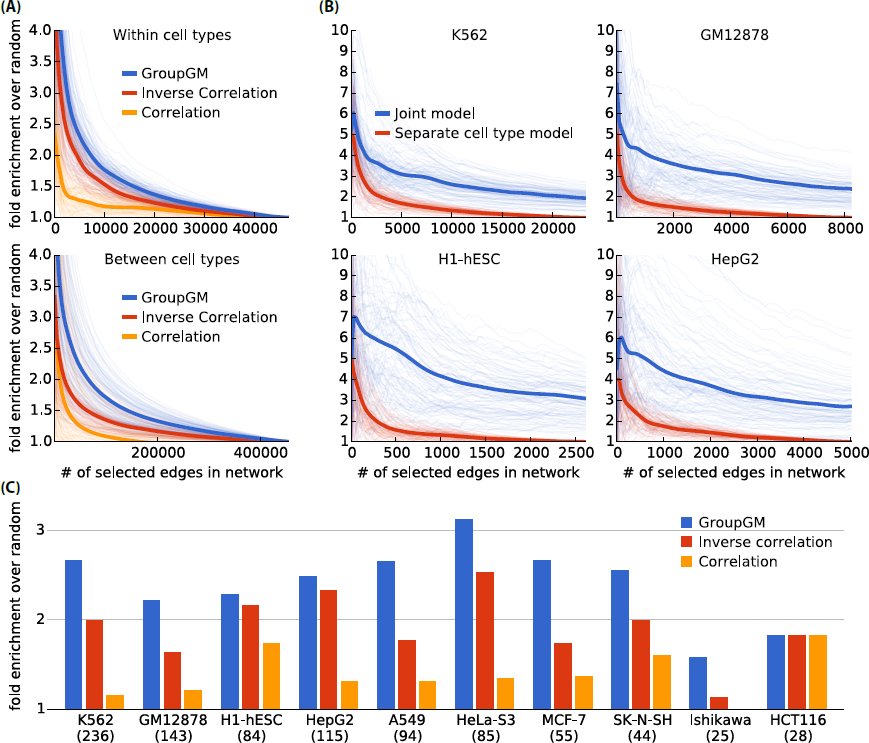
Enrichment of BioGRID-supported edges between transcription factors in networks estimated by correlation (yellow), inverse correlation (red), and GroupGM (blue). In the top plots, light lines represent bootstrap resampling variability, and dark lines represent average performance over all re-sampled networks. **(A)** Enrichment against evaluated number of network edges. *Top*: excluding edges between different cell types. *Bottom*: only including edges between different cell types. **(B)** Enrichment against evaluated number of network edges. A comparison between a joint model and a cell type specific model is repeated for four different cell types. **(C)** Enrichment within cell types that have 25 supported edges or more, where the network density was set to match number of BioGRID supported edges in each cell type. Beneath each cell type name is the number of datasets in that cell type.

**Figure 4:**
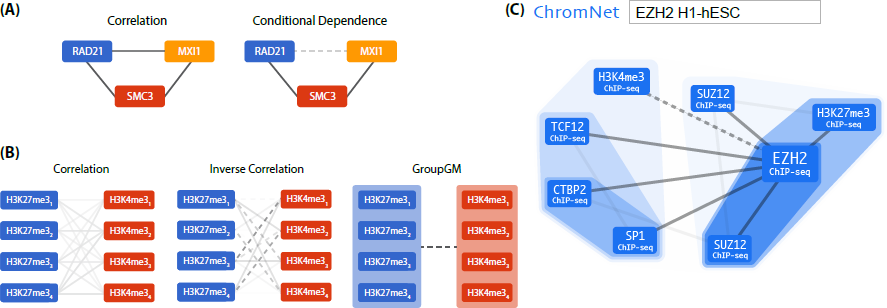
**(A)** *Left*: RAD21, MXI1, and SMC3 all co-localize with one another, suggesting they may all interact. *Right*: ChromNet reveals that the co-localization of RAD21 and MXI1 was largely mediated by the presence of SMC3. **(B)** GroupGM overcomes edge instability between tight clusters of H3K27me3 (blue) and H3K4me3 (red) modification datasets in H7-hESC at different differentiation time points. Edge darkness indicates connection strength. Dashed lines indicate negative interactions. We have removed within-group edges for clarity. *Left*: Correlation. *Middle*: Inverse correlation. *Right*: GroupGM. **(C)** The part of the ChromNet network that interacts with “EZH2 in H1-hESC” embryonic stem cells. This is a screen capture from the web interface with a search for “EZH2 H1-hESC”. Shaded regions represent GroupGM hierarchical clustering. Darker regions represent tighter clusters. Dashed lines represent negative associations. We set the edge threshold to capture the six strongest edges connected to EZH2.

To assess the variability of the enrichment estimate, we performed bootstrap re-sampling of regulatory factor targets (Figure 3A and B, light curves). All datasets with the same factor are sampled together, leading to a conservative (high) estimated variability (Methods). GroupGM showed a statistically significant improvement over both correlation (*P* = 0.0004) and inverse correlation (*P* = 0.0036) for edges within cell types (Supplementary Figure 8).

To assess variability over cell types, we estimated enrichment separately for each cell type with 25 or more BioGRID-supported edges. In each cell type, we identified the number *N* of BioGRID-supported edges in that cell type. Then, we calculated the enrichment for BioGRID-supported edges among the top *N* edges in that cell type (Figure 3C). GroupGM performed consistently better than correlation or inverse correlation in individual cell types (Supplementary Figure 9).

We also generated a simulated data set meant to mirror the characteristics of real ChIP-seq datasets (Supplementary Figure 10). Using this simulated data, we found a similar relative performance of the methods, with GroupGM recovering the most true network edges (Supplementary Figure 11; Methods).

To assess how well a joint model can recover relationships between factors measured in different cell types, we checked edges between different cell types for enrichment in known protein-protein interactions (Figure 3A bottom). The GroupGM network showed a clear enrichment for known interactions above random (*P* = 0.0095), and also out-performed inverse correlation (*P* = 0.0174) and correlation (*P* = 0.0282) (Supplementary Figure 12; Methods). This implies that information about many physical protein interactions can be recovered even from datasets in different cell types.

### Comparison between a joint model of all cell types and cell type specific models

Integrating ChIP-seq datasets from multiple cell types into a single network model provides the following three advantages. First, we can see high-level patterns in the joint chromatin network that would not otherwise be visible. Second, a joint model allows the direct comparison of cell type specific sub-networks because factors are conditioned on the same global set of ChIP-seq datasets across all cell types. Finally, a dataset for a regulatory factor in one cell type can serve as a proxy for a missing dataset for that factor in another cell type, if the factor’s localization in the genome is conserved between the cell types (Supplementary Figure 13). This greatly expands potential chromatin network edges to include the union of regulatory factors measured in any cell type. This global network contains both conserved and cell type specific sub-networks, and proves useful in analyzing data from ENCODE, which only measured a few factors in some cell types.

To directly compare a joint model across all cell types with cell type specific models for each cell type separately, we focused on the four best characterized ENCODE cell types and compared enrichment of BioGRID-supported edges (Figure 3B). By varying the number of edges in the networks we find that the joint model consistently finds interactions with higher fold enrichment for known interactions. In addition, a joint model also identifies more unique BioGRID supported protein-protein interactions than cell type specific models (Supplementary Figure 14).

We show as well that the large increase in potential edges from a joint model does not introduce spurious associations among edges within a cell type. When we excluded all cross cell type edges from the joint model, the joint model still marginally out-performs cell type specific models (*P* = 0.0672; Supplementary Figure 15).

### An example of the importance of conditional dependence: SMC3 separates RAD21 and MXI1

A specific example illustrates how conditional dependence reveals experimentally-supported direct interactions better than pairwise correlation (Figure 4A). In the correlation network among RAD21, SMC3, and MXI1, the three factors were tightly connected with one another in HeLa-S3 cervical carcinoma cells. The conditional dependence network, however, separated RAD21 and MXI1. This separation arose from the ability of SMC3 to explain away the correlation between RAD21 and MXI1. The factor pairs left connected in the conditional dependence network, RAD21-SMC3 and SMC3-MXI1, have physical interactions described in BioGRID [39, 22]. BioGRID lacks any direct connection between RAD21 and MXI1. Panigrahi et al. discovered more than 200 RAD21 interactors using yeast two-hybrid screening, immunoprecipitation-coupled mass spectrometry, and affinity pull-down assays [48]. They did not identify a RAD21-MXI1 interaction, which implies that RAD21 may not directly interact with MXI1.

In order to focus on the comparison between conditional dependence and correlation, we have not displayed the group which joins SMC3 and RAD21. This grouping reflects their common role in the cohesion complex and is present in many cell types. We also note that Figure 4A is only a small part of the full ChromNet network and considering more factors reveals additional relationships that involve CTCF and ZNF143, which is consistent with prior knowledge [74] (Supplementary Figure 16).

### An example of the importance of group dependency: recovering a connection between H3K27me3 and H3K4me3

Another specific example shows how GroupGM mitigates the effect of redundancy on conventional conditional dependence models. We examined edges between multiple H3K27me3 and H3K4me3 datasets from H7-hESC embryonic stem cells, collected at different time points in differentiation [47]. H3K27me3 is a repressive mark and H3K4me3 is an activating mark. Since the datasets observe discrete portions of the differentiation process, one should not average them or pick a reference dataset arbitrarily. However, the H3K27me3 datasets are correlated highly enough with one another to form a group, and so are the four H3K4me3 datasets. This implies that conventional conditional dependence methods would identify edges between the two histone marks incorrectly.

Edges estimated using correlation indicate that the datasets targeting H3K27me3 and those targeting H3K4me3 are positively correlated. However, H3K27me3 is associated with repressed genomic regions while H3K4me3 is associated with actively transcribed regions [73]. A minority of promoters in embryonic stem cells are "bivalently" marked, but this should not lead to an overall positive association [5, 73]. In fact, most ChIP-seq datasets are positively correlated with each other (Supplementary Figure 13), which is induced by mappability and many regions that are transcriptionally silent or active. Resolving this problem by removing some of these regions is likely to fail, because it is not clear based on what criteria we need to exclude regions. In the conditional dependence models, such as inverse correlation and group graphical model, by conditioning on many other variables, these global confounding effects are naturally removed. Edges estimated by inverse correlation account for these confounders but become weak and unstable showing a mixture of positive and negative associations (Figure 4B, middle). By allowing group edges, GroupGM has power to recover the negative association between H3K27me3 and H3K4me3 (Figure 4B, right), which is consistent with prior knowledge.

### An example of learning genomic context: ZNF143 mediates the conditional dependence relationship between CTCF and SIX5

Many relationships between regulatory factors only occur in a particular genomic context. This raises the question of how, or whether, this context specificity is encoded in ChromNet. We can gain insight into this by considering what it means for one factor to mediate the relationship between two other factors, such as A mediating the relationship between C and D in Figure 1A. When this occurs it means that the connection between C and D can be explained away by their co-occurrence with A. In other words, A is the context in which the relationship between C and D occurs.

A practical example of this is found in the relationships between SIX5, ZNF143, and CTCF in the K562 cell type. Simple correlation connects all three factors together with positive edges, but GroupGM shows that ZNF143 actually mediates the relationship between SIX5 and CTCF (Supplementary Figure 5; Supplementary Figure 6). This means that the association of SIX5 with CTCF primarily occurs in the presence of ZNF143, the CTCF-SIX5 relationship is context specific and ZNF143 is the context. More generally when an association between two factors, C and D, is specific to a certain genomic context and that context is well represented by a third factor, A, then A will mediate C and D. This gives the connections C – A – D in the conditional dependence network; thus context-specific relationships, such as the relationship between CTCF and SIX5 in the presence of ZNF143, are captured in a GroupGM network, if all three factors are present.

It is important to understand the genomic context in which any given edge occurs regardless of whether that context is well represented by another factor in the network. Even if A is not observed we want to be able to infer the genomic context of the interaction between C and D. To address this need we designed an efficient method to label every genomic position with its influence on a group network edge (Methods). Using CTCF–ZNF143–SIX5 as an example, we removed all ZNF143 experiments from ChromNet and then computed the genomic context of the edge between CTCF and SIX5. To validate this genomic context, we took the top 1,000 bins (1,000,000 bp) and intersected them with the top 1,000 bins from all other experiments in K562, including ZNF143. Even though ZNF143 was not present in the model and ZNF143 datasets were not used when inferring the genomic context, it had the highest overlap of any experiment with the context driving the CTCF-SIX5 edge, even higher than the CTCF and SIX5 experiments themselves (Supplementary Figure 17).

### An example of network accuracy: recovered interactions with EZH2 in H1-hESC recapitulate known functions

As an example illustrating the utility of ChromNet in revealing the potential interactors of a specific regulatory factor we examined a small portion of the network associated with the well-characterized protein EZH2 (Figure 4C). We focused on the H1-hESC cell type because it had many strong EZH2 connections in ChromNet.

Examining connections to EZH2 in H1-hESC highlighted several known interactions, which we discuss in decreasing order of edge strength. The strongest connection is from H3K27me3, and EZH2 is a methyltransferase involved in H3K27me3 maintenance [1]. The next strongest connections are to SUZ12, which is an essential part of the Polycomb repressive complex 2 (PRC2), and is required for EZH2’s methyltransferase activity [8, 14]. The next connection to CTBP2 is supported by this co-repressor’s possible role in deacetylation of H3K27 in preparation for PRC2 mediated methylation [29]. H3K4me3 is well known to be present in active regions of the genome, so a negative relationship with EZH2 (represented by a dashed line) that deposits the repressive H3K27me3 mark is expected. SP1 is a potentially novel connection with EZH2, while the final linked factor, TCF12, co-immunoprecipitates with EZH2 [37]. In summary, most of the strongest interactions with EZH2 have support in the literature. We found this mixture of positive controls and potential novel connections in many parts of the network.

### An example of cross cell type comparison: enhancer associated regulatory factors

Learning a conditional dependence network for all ENCODE cell types allows the comparison of within cell type connections across different cell types. Active enhancers are known to be flanked by histones marked by a combination of H3K27ac and H3K4me1 [58]. So to quantify how strongly different transcription factors associate with active enhancers in different cell types we calculated the sum of the group edges between each regulatory factor (except histone marks) and H3K27ac and H3K4me1 measured in the that cell type. This provides a score for each factor in each cell type.

Seven ENCODE cell types with 20 or more datasets contain both H3K27ac and H3K4me1, while also containing EP300, which is known to bind active enhancers [58]. We focused on these seven cell types and ranked the factors in each cell type by their association with H3K27ac and H3K4me1. Supplementary Table 3 lists the top 10 factors in each cell type most associated with active enhancers. EP300 can be considered a validation for the list and is highly ranked in all seven cell types (*P* < 10^−5^). Interestingly, even more highly ranked than EP300 is POLR2A. This association is likely because active enhancers are in close proximity to active transcription start sites in promoters in a 3D space, due to the looping mechanisms for enhancer-promoter communication. The influence that 3D conformation can have on measures of co-localization in the genome is important to bear in mind when analyzing ChromNet edges. Other factors that are consistently associated with enhancers across cell types are shown in red, while cell type specific associations are in black.

### An example of a novel protein interaction: experimental validation of an interaction between MYC and HCFC1

The c-MYC (MYC) transcription factor is frequently deregulated in a large number and wide variety of cancers [42, 52]. It heterodimerizes with its partner protein MAX to bind an estimated 10-15% of the genome to regulate the gene expression programs of many biological processes, including cell growth, cell cycle progression, and oncogenesis [42, 52, 6]. The mechanisms by which MYC regulates these specific biological and oncogenic outcomes are not well understood. Interactions with additional co-regulators are thought to modulate MYC’s binding specificity and transcriptional activity [23, 64]; however, only a few MYC interactors have been evaluated on a genome-wide level. Analysis of the large number of ENCODE ChIP-seq datasets can therefore further elucidate MYC interactions at the chromatin level.

ChromNet showed that MAX is the strongest interactor of MYC across multiple cell types (Supplementary Table 2), highlighting the ubiquitous nature of this interaction. Top-scoring ChromNet connections also included other known MYC interactors, like components of the RNA polymerase II complex such as POLR2Aand chromatin-modifying proteins such as EP300 (Supplementary Table 2). This shows how ChromNet analysis of transcription factors and co-regulators can help better understand transcription factor complexes.

In addition to the known interactors above, ChromNet also revealed previously uncharacterized, high-scoring interactions, including the transcriptional regulator Host Cell Factor C1 (HCFC1; Supplementary Table 2). HCFC1 binds largely to active promoters [43] and is involved in biological processes, such as cell cycle progression [51, 55] and oncogenesis [50, 13, 53]. This further supports its possible role as an interactor of MYC in regulating these activities. To validate the novel MYC-HCFC1 interaction, we performed a proximity ligation assay (PLA) in MCF10A mammary epithelial cells. This technique detects endogenous protein-protein interactions in intact cells [59] and has been used to validate novel interactors of MYC [19]. When two proteins that are probed with specific antibodies are within close proximity of each other, fluorescence signals are produced that are measured and quantified using fluorescence microscopy. We saw only background fluorescence when incubating with antibody against MYC (Figure 5A, top) or HCFC1 (Figure 5A, middle) alone. Incubation with both MYC and HCFC1 antibodies yielded a significant increase in fluorescence signal in the nuclear compartment (Figure 5A, bottom; Figure 5B). This suggests that MYC and HCFC1 interact in the nucleus.

**Figure 5:**
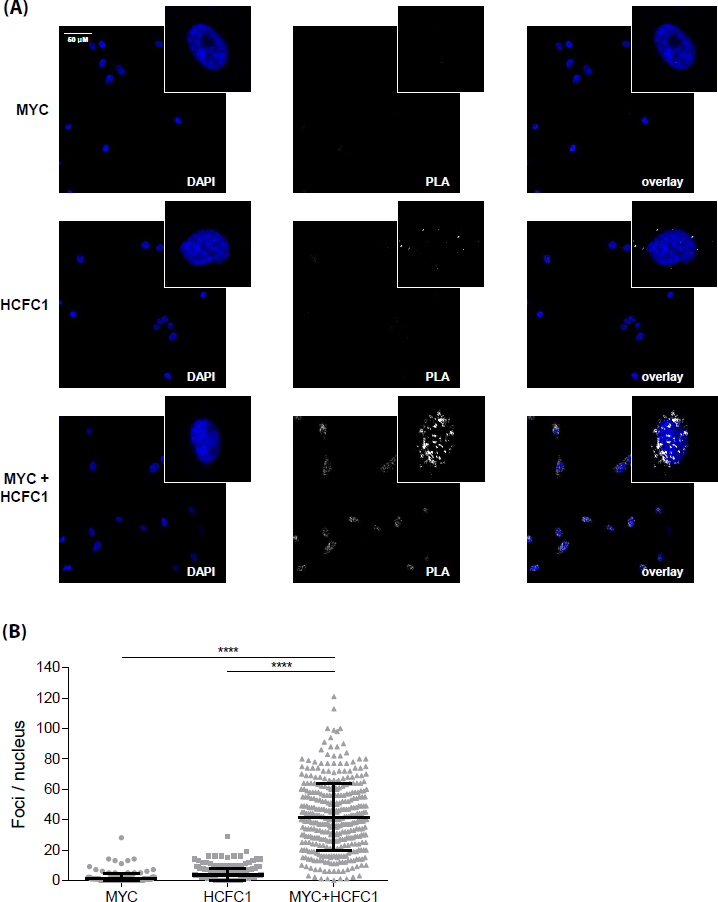
**(A)** Proximity ligation assay showing MYC and HCFC1 interaction in the nucleus. Representative micrographs show DAPI nuclear staining (left), proximity ligation signal (middle), and overlay (right) at 20× magnification, with insets at 100 × magnification. *Top*: Cells probed with MYC antibody alone. *Middle*: Cells probed with HCFC1 antibody alone. *Bottom*: Cells probed with both antibodies. **(B)** Proximity ligation assay signal quantified as number of foci per nucleus, with 254, 293, and 381 nuclei quantified for the MYC antibody alone, the HCFC1 antibody alone, and both antibodies together conditions, respectively. Individual values (grey dots) and mean ± standard deviation black bars from three biological replicates are shown; **** *p* < 0.0001, one-way analysis of variance with Bonferroni post test. Quantifications for each independent replicate are shown in Supplementary Figure 18.

We have shown that HCFC1 may be a novel co-regulator of MYC. Future investigation will reveal the importance of HCFC1 in regulating the biological functions of MYC, such as cell cycle progression and oncogenesis. This discovery illustrates how ChromNet can suggest novel protein-protein interactions within chromatin complexes.

### Spatial embedding reveals global patterns in the human chromatin network

By integrating all ENCODE datasets from many cell types into a single network, ChromNet enables extraction of global patterns in the relationships among regulatory factors. We used multidimensional scaling [7] to embed the entire network into a 2D layout (Figure 6; Methods). In this embedding, the spatial proximity of two nodes is designed to reflect their distance in the network, where positive edges pull nodes closer together and negative edges push them father apart. Nodes for the same regulatory factor in different cell types form a cluster when that factor’s genomic position is conserved across cell types. For example, CTCF forms a clear cluster in this manner (Figure 6A). Relationships between regulatory factors are represented by their proximity in the embedding. For example, MYC and MAX nodes are located in the same region. So are CTCF and RAD21. In contrast to the joint network, relationships in individual cell type networks (Figure 1B top) are much less distinct (Supplementary Figure 19).

**Figure 6:**
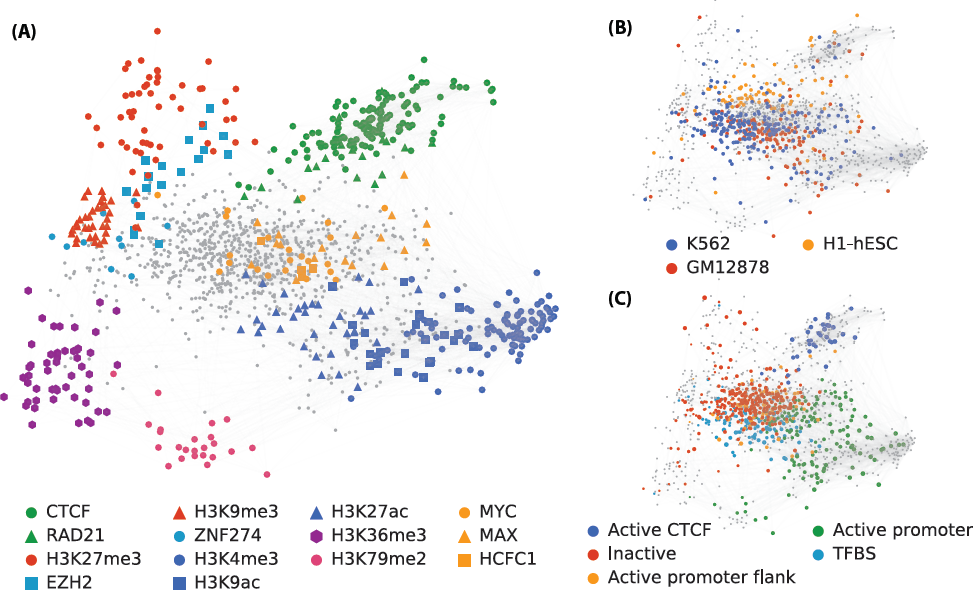
2D embedding of the entire human chromatin network estimated by ChromNet. Spatial proximity of nodes and node groups reflects the strength of their inferred connection. In three views of this embedding, we have highlighted three different aspects: **(A)** Specific regulatory factors discussed in this article. **(B)** Datasets from the three ENCODE Tier 1 cell types, showing a separation of regulatory factors by cell type. **(C)** Correlation with five Segway genome annotation labels. We only colored the datasets from cell types where the Ensembl Regulatory Build had a corresponding Segway annotation. Node size represents correlation to a label, in comparison to all other nodes assigned to that label.

Relative positions of regulatory factors in the embedded graph highlight important aspects of biology. This is especially apparent among histone marks, where there is a clear separation between activating marks such as H3K4me3 and H3K27ac on the lower right and repressive marks such as H3K27me3 and H3K9me3 on the upper left (Figure 6A). H3K9me3 and H3K27me3 are both repressive marks, but form distinct clusters because they target distinct regions of the genome. H3K27me3 marks facultative heterochromatin, thought to regulate temporary repression of gene-rich regions [28]. H3K9me3 marks constitutive heterochromatin, and acts as a more permanent repressor [32]. Between the active and repressive marks we find H3K36me3 and H3K79me2. H3K36me3 is closer to the inactive marks and is implicated in restricting the spread of H3K27me3 [69]. H3K79me2 varies with the cell cycle and is associated with replication initiation sites [18]. The relative position of histones and protein factors is also interesting. ZNF274 has been implicated in the recruitment of methyltranferases for H3K9me3 and is found nearby in the network [17]. EZH2 is involved in the deposition of H3K27me3 and is found between the H3K27me3 cluster and the rest of the network [67].

Positions of regulatory factor datasets reflect both their cell type identities and association with chromatin states. Highlighting the three Tier 1 ENCODE cell types shows a weak clustering of regulatory factor datasets by cell type (Figure 6B). K562 and GM12878 are both derived from blood cell lines and overlap more spatially with one another in the network than with H1-hESC (embryonic cells). Coloring the network by correlation with chromatin state also reveals spatial patterns. We chose five (out of seven) Segway [25, 70] annotation labels that highlight distinct areas of the network (Figure 6C), illustrating a clear separation between active and inactive regions of the genome, and that chromatin domains are reflected in the interactions of the chromatin network. Spatially embedding regulatory factor datasets using the ChromNet network simultaneously captures many important aspects of their function, such as chromatin state, cell lineage, and known factor-factor interactions.

## Discussion

### ChromNet enables understanding the chromatin network

Characterizing the chromatin network, the network of interactions among regulatory factors, is a key part of understanding gene regulation. ChromNet provides a new way to learn the chromatin network from ChIP-seq data. ChromNet addresses key problems encountered when learning a joint network of protein-protein interactions from ChIP-seq datasets, such as the need to distinguish direct from indirect regulatory factor interactions while remaining robust to dataset redundancy. ChromNet also provides an efficient method to learn the genomic context driving an edge, which allows a more comprehensive understanding of the inferred interactions. We demonstrated that ChromNet’s GroupGM network infers known protein-protein interactions in the joint chromatin network more accurately than other methods.

Unlike many previous methods, ChromNet is also efficient enough to integrate thousands of genome-wide ChIP-seq datasets into a single joint network. To our knowledge, this study represents the first construction of an interaction network from all 1,451 ENCODE ChIP-seq datasets. ChromNet already scales to the number of datasets necessary to represent all 1,400–1,900 human transcription factors [66], once such data is available.

ChromNet provides a general computational framework to identify a joint dependence network from many ChIP-seq datasets. It can build a custom joint dependence network by incorporating user-provided ChIP-seq datasets or a combination of the ENCODE ChIP-seq datasets and user-provided datasets. To allow easier exploration of regulatory factor interactions and to facilitate generation of novel hypotheses, we have created a dynamic search and visualization web interface for both the ENCODE network and networks built from custom datasets (http://chromnet.cs.washington.edu). By building a large model and allowing easy inspection of small sub-networks, ChromNet combines a large-scale conditional dependence model with practical accessibility.

To demonstrate ChromNet’s ability to reveal novel regulatory factor interactions, we experimentally validated the interaction between the MYC and HCFC1 proteins. The biological functions of the MYC oncoprotein are complex and dependent on its protein-protein interactions. Uncovering these interactions will provide insights into MYC transcriptional complexes involved in the oncogenic process and may also reveal potential targets for anti-cancer therapies. While this manuscript was under review, the MYC-HCFC1 interaction was independently described by Thomas et al. (2015), further strengthening our validation of the interaction discovered through ChromNet and establishing HCFC1 as a bona fide interactor of MYC. Through ChromNet, we identified HCFC1 as a novel interactor of MYC that may be involved in regulating biological and oncogenic functions of MYC.

### Future directions

We envision several future extensions to the approach described in this article. First, while we have demonstrated the utility of applying ChromNet to ChIP-seq data alone, we plan to incorporate other data types into the network. RNA-seq expression datasets could resolve regulatory factor relationships that occur as a consequence of mutual involvement in gene expression. Incorporating feature annotations such as gene models could highlight direct interactions between factors and genomic regions of interest. The human genome’s billions of base pairs provide a large sample size that allows joint comparisons of many genome-wide signals in a single model. Robust conditional dependence networks provide a benefit that is likely not limited to ChIP-seq data.

Second, we plan to consider relationships between regulatory factors at genomic position offsets. Here, we considered only co-occurrence relationships within the same 1,000 bp region. To model positional ordering constraints, we can also consider relationships between a factor in one region and another factor in an adjacent or nearby region. This would allow us to learn phenomena such as promoter–associated factors preceding gene-body–associated factors.

Third, just as the co-occurrence of different regulatory factors has been used to automatically annotate the genome, variations in the chromatin network at different positions may also prove useful to annotate functional genomic regions. This would also provide insight into the biological mechanisms behind specific regulatory factor interactions and the chromatin states in which they occur.

## Methods

### Data processing

ENCODE has the largest collection of high-quality ChIP-seq datasets [11], and continues depositing new datasets. ENCODE has processed many ChIP-seq datasets through a uniform pipeline. However, we reprocessed all the datasets from raw ChIP-seq reads (Figure 2) for two reasons. First, this allowed us to incorporate datasets not yet through ENCODE’s uniform pipeline. Second, specifying our own pipeline makes it easier to process external users’ data in an identical way. This facilitates adding ChIP-seq datasets that are not from the ENCODE project to the ChromNet network.

We aligned reads from 3,574 FASTQ files to GRCh38/hg38 [20] using Bowtie2 [33]. We grouped BAM files by dataset using metadata from the ENCODE web site [16]. Then, we pooled and processed BAM files using a custom binning method that counts the number read starts in each of 3,209,287 1,000 bp bins covering all contigs in GRCh38/hg38.

Binning all count datasets yielded a 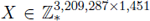 count-valued *data matrix*. Each bin has a corresponding row in the matrix. We interpreted each of the 3,209,287 rows as a sample from a set 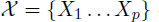 of *p* = 1,451 count-valued random variables representing occupancy of each regulatory factor at a given position. Using this interpretation, we computed a sample correlation matrix 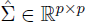 among the standardized variables in 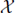.

To create the *correlation network*, we set the weight of every edge between two datasets *i* and *j* equal to the corresponding entry 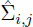 in the sample correlation matrix. This captures the pairwise linear dependence between two datasets (Figure 1A; bottom left).

### Generation of simulated data

A large-scale simulated dataset was generated to validate the ability of ChromNet to recover interactions from raw count data. Representing the conditional dependence between large numbers of count variables for the purpose of simulation is not trivial. It is important that the model is not overly simplistic, but also still interpretable. Here we use a multivariate Gaussian distribution to represent the means of marginal Poisson distributions, thresholding the values when they fall below zero, and adding additional negative binomial distributed noise to represent random reads unrelated to protein localization.

In this model the count for a track *j*, *c_j_*, at a given position is described by

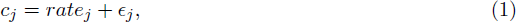

where *rate_j_ ∼ Poisson(max(0, s_j_))* and *є_j_ ∼ NegativeBinomial(r,p)*. The signal follows a thresholded normal distribution *s_j_ ∼ max(N(µ,Σ*),0). The background noise (*r*= 25,*p*= 0.9) and the parameters of the normal distribution are all fixed during the course of the simulation. 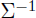 represents the structure of the underlying conditional dependence network.

The inverse covariance matrix of the simulated data 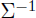 was randomly generated with a sparsity of 10%. In addition, datasets were grouped into complexes of size one (60%), two (20%), or three (20%) to represent the type of close coupling observed in real data among some factors. The correlation within complexes was set to between 0.8 to 0.9 to match the magnitudes of high correlations observed in real data (Supplementary Figure 2). A total of 80 complexes were simulated across 200,000 positional samples. This results in 126 experiments and 200,000 samples.

To better model the complexities found in real data sets we added dependence between nearby samples by convolving the signal data with a [0.05, 0.9, 0.05] kernel before adding noise. This caused nearby bins to be more similar to each other and thus the samples are not identically independently distributed. Since larger correlations between regions of the genome are also present due to batch effects or other confounding factors we added one of 8 different batch effect tracks to each of the 126 datasets.

The resulting marginal count distributions from this model are visually similar to those observed in real data (Supplementary Figure 10). Because we based the correlations between datasets on a (largely transformed) multivariate normal distribution we can treat datasets connected in the underlying generative model as truth and seek to recover them using a variety of methods. The results of this analysis are shown in Supplementary Figure 11, which is consistent with Figure 3, where the group graphical model performs better than alternative approaches including correlation, inverse correlation and partial correlation.

### Efficient estimation of conditional dependence from count data

Given datasets drawn from a set 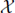 of count-valued random variables, learning an exact joint model that captures the dependency structure of these datasets could be challenging. Although there are a variety of multivariate count distributions, all are either overly restrictive or challenging to estimate for large numbers of variables [68].

A common alternative is to use a multivariate Gaussian distribution and some type of transform on the marginals to make them more Gaussian such as *sqrt* or *asinh*. Since count data is often heteroscedastic, where variance increases with higher counts, these transforms squash higher values, making the distribution more symmetric. This causes the least squares error term to focus less on high valued samples and proportionately more on lower values. Interestingly, for ChIP-seq datasets this is not desirable because higher values are more likely to represent strong signal while lower values are more likely driven by noise.

Because of its efficiency and interpretability we used a multivariate Gaussian approximation to the count data for ChromNet. We also chose to use untransformed raw read counts in the model. This choice was based on observing a clear decrease in performance when using transforms designed to mitigate heteroscedasticity (Supplementary Figure 1).

An additional benefit of using a multivariate Gaussian is that it can also serve as a reasonable approximation to a Markov random field distribution. This allows for the comparison with other methods designed to work strictly with binary data (Supplementary Figure 20; Supplementary Figure 21; Supplementary Note 3).

To create the *inverse correlation network* (Figure 3), we began by inverting the sample correlation matrix 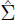 to get an inverse sample correlation matrix 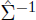 [40, 35]. We then set the weight of every edge between two datasets *i*and *j*equal to the corresponding entry 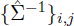. This inverse correlation network captures the pairwise linear dependence between two datasets when conditioned on all other variables in the network.

It should be noted that partial correlation is very similar to inverse correlation and has been used before by Lasserre et al. to effectively model connections between histone marks from human ChIP-seq data (using rank-transformed data from gene start sites) [34]. The matrix of partial correlations, *P*, is a renormalization of inverse correlation 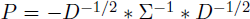 where *D* is the diagonal matrix of 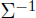. A direct application of partial correlation to all ENCODE data suffers from the same issues as inverse correlation, performing slightly worse in the recovery of known protein-protein interactions (Supplementary Figure 3). We chose to use inverse correlation as the foundation of the group graphical model (GroupGM) because the proof that GroupGM recovers the correct edge weights in the presence of near perfect redundancy does not hold when applied the to partial correlation matrix (Supplementary Note 2).

One additional concern when applying a Gaussian graphical model to ChIP-seq data is that the values at each 1,000bp bin in the genome are not independent of each other. Fortunately, while this may reduce the power of the model (i.e. it will need more samples), it does not bias the model. This is because the edges of a Gaussian graphical model can be interpreted in terms of linear regression coefficients. Standard linear regression coefficients are unbiased even when samples are not statistically independent when the data follows a linear relationship. To validate this on ChIP-seq data, and to confirm that any loss of power is unimportant we evenly subsampled the data at progressively larger intervals. We found that performance when recovering known protein-protein interactions does not degrade until we subsample 100-fold (Supplementary Figure 22).

### Group graphical model

To create the *Group Graphical Model (GroupGM) network*, we began with the inverse correlation matrix created above. We extended the idea of pairwise relationships to groups of datasets by considering a set 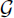 of *q* groups chosen by hierarchical clustering (see below). This effectively allows edges to express relationships between groups of variables. We let 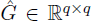 represent pairwise interaction strengths between all groups in the model. For any two groups *i* and *j* in the model their weight is given by the sum of entries between them in the inverse correlation matrix (Figure 1C).

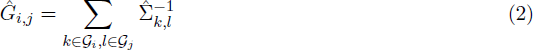

We prove that Equation 2 correctly maintains the original edge magnitude in the case of redundancy (Supplementary Note 2).

To select the set 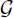 of groups, we used complete-linkage hierarchical agglomerative clustering of the correlation matrix [24]. This clustering method starts by merging the two groups with the smallest maximum correlation distance between their datasets, then continues recursively until all groups have been merged. The use of hierarchical clustering eliminates the need to chose a fixed arbitrary number of clusters in advance. From the clustering results we chose all the leaf and internal nodes from the clustering algorithm as groups 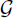. Then, *G* became a *q*×*q* matrix filled according to Equation 2, where *q* = 2*p*– 1 (the total number of internal and leaf nodes). This method avoids comparing all possible subsets of datasets, which would make calculating *G* prohibitively expensive. Since groups with low correlation are less likely to cause the collinearity problem (Figure 1C), we only consider groups with a correlation greater than 0.8 which captures 53% of all the multi-factor groups formed by the hierarchical clustering (Supplementary Figure 2).

Since GroupGM uses the cluster assignments to mitigate strong redundancy, clustering accuracy is most important for tightly correlated datasets. When two datasets are highly correlated, it is important to group them together to mitigate the outcome of correlated datasets in network inference. When two datasets are only mildly correlated, the effects of their redundancy will also be mild, so it is less important to group them together. Hierarchical clustering is an attractive choice because it starts by creating groups among the most correlated datasets.

### Computing the genomic context that drives a network edge

The conditional dependence relationships represented by an edge in ChromNet can occur primarily in certain genomic regions. Here we seek to identify what parts of the genome (i.e. samples) drove the creation of an edge in ChromNet. Understanding what positions in the genome caused ChromNet to estimate a network edge provides insight into the genomic regions driving the relationship.

The most natural way to define the influence of a genomic position (i.e. sample) on an edge is as the difference in edge value between when we observe a position and when we do not observe a position in the genome. If implemented directly this could easily become computationally intractable since it involves relearning the entire model for every position in the genome. For a highly optimized implementation on 16 cores, computing the correlation matrix takes approximately two minutes, which would lead to a run-time of over 12 years for 3,209,287 binned genomic positions. This can be sped up dramatically by using rank-1 matrix updates to avoid recalculating most of the correlation matrix. This results in a much faster method, where the slowest step is the inversion of the correlation matrix. However, computing this inversion for each genomic sample still leads to over four days of computation on recent high performance servers. Pre-computing this information is also undesirable since it would create 54TB of largely incompressible data for all group edges. Below we show that for the ChromNet model, the calculation of s genomic position’s impact on an edge can be made extremely efficient. The ideas are similar to those used in efficient leave-one-out cross validation implementations for linear models.

Removing a genomic position and computing the new inverse correlation matrix can be written in terms of a rank-1 update and the inverse correlation matrix before the position (sample) is removed. This equation holds under the assumption that removing the sample does not change the mean of the data. Let Σ be the correlation matrix of all the data, and 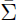 be the correlation matrix with the sample removed. Let *u* be the column vector representing the sample to be removed (already mean centered). Letting *D* be a normalizing diagonal matrix 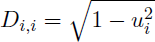 we get:

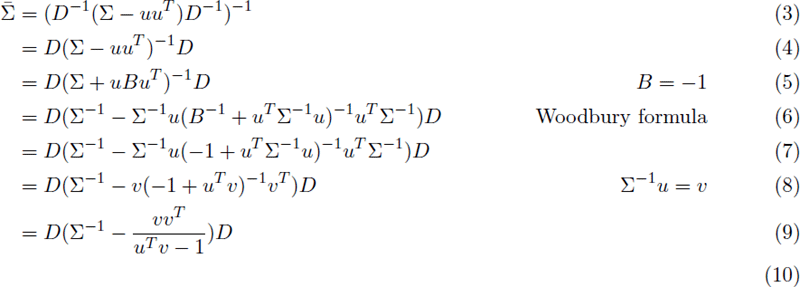

Included in the ChromNet software release is an optimized implementation utilizing the above inverse rank-1 update formulation. It can solve 40,000 model updates to the full joint chromatin network per second, which leads to a run-time of just over one minute for a single group edge over the human genome. The output is the effect each genomic position has on an edge when that position is added to the dataset. This information can be used to examine the highest impact positions and determine the genomic context driving an edge (Supplementary Figure 17).

### Visualization of the hierarchical chromatin network

To enable exploration of the chromatin network, we built an interactive visualization tool (http://chromnet.cs.washington.edu). This tool displays the nodes and edges of the chromatin network using a real time force model (Figure 4C). The tool’s responsive interface lets users control which nodes and edges it displays. It immediately changes its display after a user types a search term to restrict displayed nodes. It also immediately changes its display when a user moves a slider that controls the minimum strength of a displayed edge. Our visualization tool facilitates exploring the chromatin network without excessive visual distraction.

The ChromNet visualization tool displays hierarchical groups from GroupGM by shading areas that enclose a group’s members. It shades these areas with some amount of transparency. It displays the strongest groups with the highest opacity. The parents of two connected groups in the GroupGM hierarchy are themselves very likely connected. Therefore, for clarity we hide redundant parental edges.

To find a reasonable lower bound for the user-defined strength threshold, we examined the relationship between edge magnitude and known physical interactions. Within cell type edges from all cell types were sorted by magnitude and then binned. For each bin we computed the number of edges matching low throughput physical interactions in BioGRID and plotted how this varied over the bins. This enrichment curve suggested a lower bound of 0.2 to capture only edges enriched for known interactions (Supplementary Figure 23).

### Fold enrichment reflects both type I and type II error rates

The fold enrichment is a single quantity that captures the effects of both type I and type II error rates. This can be seen from the definition of fold enrichment:

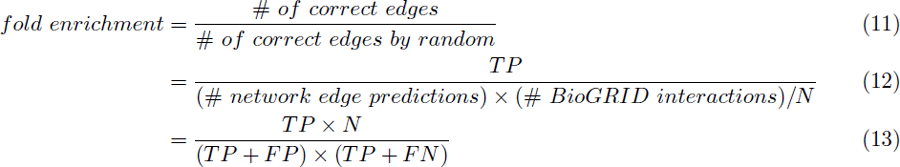

where *N* is the total number of possible edges, and *TP, FP*, and *FN* refer to the number of true positives, false positives, and false negatives, respectively. The fold enrichment is inversely proportional to the number of false positives (type I error) and number of false negatives (type II error). The type I error rate is equal to (type I error) / (total number of BioGRID interactions), and type II error rate is equal to (type II error) / (total number of interactions - number of BioGRID interactions). Since the denominators of the type I and type II error rates are fixed numbers, we can say that the fold enrichment is inversely proportional to the type I and type II error rates.

### A conservative bootstrap estimate of protein-protein interaction enrichment variability

We estimated the variability of enrichment for known protein-protein interactions in the chromatin network (Figure 3) using bootstrap re-sampling over regulatory factors. We performed re-sampling over regulatory factors, and not over edges or individual datasets, because valid bootstrap re-sampling assumes independent and identically distributed samples. If we had re-sampled over the edges, we would have estimated a much smaller variability. This is because edges do not vary independently, and changes in a single dataset can affect all edges connected to that dataset. Variation specific to a single regulatory factor would affect all datasets measuring that factor. Those individual datasets, therefore, lack the independence assumed by the bootstrap sampling.

Under a regulatory factor bootstrap, we might sample a widely-measured regulatory factor a number of times. For example, ChromNet contains 130 CTCF datasets. Every time we sample CTCF, we add all 130 of these columns (where a column represents a variable in the data matrix *X*) to the bootstrap data matrix. Adding many datasets in unison greatly increases variability in the re-sampled data matrix. This yields conservative high variability estimates, ensuring that enrichment performance is not solely due to a few commonly measured factors. Using these bootstrap samples, we compared the area under the enrichment rank curves (Figure 3A,B) between methods. The statistical significance of GroupGM’s improvement was quantified as the fraction of bootstrap samples where GroupGM outperformed the other methods (Supplementary Figure 8; Supplementary Figure 12).

### Proximity ligation assay

We seeded 2.5 × 10^4^ MCF10A cells (a kind gift from S. Muthuswamy, Princess Margaret Cancer Centre) onto glass cover slips. After one day, we fixed cells in 2% paraformaldehyde, permeabilized the cells, and blocked them with bovine serum albumin. We then incubated the cells overnight with a mouse monoclonal antibody against MYC (1:25; C-33, Santa Cruz Biotechnology, Dallas, TX) and a rabbit polyclonal antibody against HCFC1 (1:50; A301-400, Bethyl Laboratories, Montgomery, TX) overnight. Then, we incubated cells with Duolink In Situ PLA anti-mouse MINUS and anti-rabbit PLUS probes (Sigma-Aldrich, St. Louis, MO). We processed cells using Duolink In Situ Detection Reagents Red following manufacturer’s instructions (Sigma-Aldrich, St. Louis, MO). We imaged six fields of view per slide with a LSM700 confocal fluorescence microscope (Zeiss, Oberkochen, Germany). We unbiasedly quantified proximity ligation assay signal per nucleus (as defined by DAPI staining) using the software ImageJ [57].

### Embedding the full chromatin network into a single plot

Embedding a graph into a space involves defining distances between all nodes in the graph. Because the GroupGM is inherently multi-scale, we sought a distance metric that accurately represented forces between individual nodes, and between all possible node groupings. In GroupGM, the edge weights between two groups are sums of the conditional dependence weights between all the individual datasets of those groups. A common method of computing graph distances that accounts for the total effect of all edges between two groups is the resistance distance [30]. The name is derived from an interpretation of the distance as the electrical resistance between two nodes in the graph where edges are viewed as wires. This can be computed as:

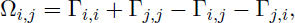

where Γ is the inverse of the graph Laplacian.

While at first glance the resistance distance may seem like an arbitrary metric to use for node distances, upon closer inspection we find striking parallels between it and Gaussian graphical models. First note that the weighted graph laplacian [45], *L*, is defined as:

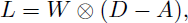

where *D* is a diagonal matrix of edge degrees, *A* is the binary adjacency matrix of the graph, *W* is a matrix of positive edge weights, and 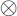 represents element-wise multiplication. A general Gaussian graphical model has a complete graph so *A* will be all ones, and *D* will be constant on the diagonal, the edge weights will be symmetric and can be positive or negative. Positive edge weights will lead to negative off diagonal entries in *L*, just as positive connections in the GGM will lead to negative off-diagonal entries in 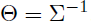. So by allowing *W* to contain negative entries we can view 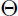 as a type of graph laplacian.

Viewing 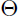 as a type of graph Laplacian allows us to compute the resistance distance by setting 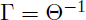. Simplifying gives the following

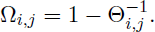

So the resistance distance is just a constant offset of the correlation matrix of the network. This means that if we are trying to compute distances between nodes in a graph represented by the inverse correlation matrix, correlation is a very natural distance measure. We note however that unlike the original data correlation matrix this matrix is computed from the inverse of the edge weights matrix. This causes a difference because we threshold small edge values that are likely to only represent noise. We chose this threshold to maximize the visual clarity of the network, which lead to a threshold of 0.01.

We overlaid chromatin state annotation on the graph embedding by computing the correlation between each dataset and each Segway [25] region from the Ensembl Regulatory Build for GRCh38/hg38 [70]. We drew a separate network labeling for each region by sizing each dataset node by its correlation with that Segway region. We normalized the size of the largest node in each network to a constant value and overlaid three of these network colorings (Figure 6C).

## Acknowledgements

We would like to acknowledge William S. Noble, Zhiping Weng, R. David Hawkins, W. Larry Ruzzo, and Maxwell W. Libbrecht for their helpful feedback during the development of ChromNet.

This work was supported by a National Science Foundation (NSF) Graduate Research Fellowship (DGE-1256082) to S.M.L.; NSF (DBI-1355899) to S.I.L.; Natural Sciences and Engineering Research Council of Canada (RGPIN-2015-03948) to M.M.H.; Canada Research Chair in Molecular Oncology to L.Z.P., Canadian Institute for Health Research (MOP-275788) to L.Z.P. and B.R; and a Canadian Breast Cancer Foundation Ontario Region Doctoral Fellowship to W.B.T. Cloud computing resources for this research were generously provided by Google.

The authors declare that they have no competing interests.

